# A single kindred across centuries: patrilineal continuity and a female-mediated increase in the local genetic component of the Lake Onega area at the Kondopoga cemetery

**DOI:** 10.64898/2026.11.13.766448

**Authors:** D.A. Vashunina, M.V. Leonova, I.K. Reshetova, A.V. Nikitina, M.V Dobrovolskaya, K.V. Zhur, N.A. Makarov, E.B. Prokhortchouk

**Affiliations:** Federal Research Center for Biotechnology, Russian Academy of Sciences, Moscow, Russian Federation; Institute of Archeology, Russian Academy of Sciences, Moscow, Russian Federation; Sirius University, Sochi, Russian Federation; Pirogov Russian National Research Medical University, Moscow, Russia

## Abstract

We report the first genome-wide ancient DNA data from a medieval population in the territory of the modern Republic of Karelia, Russia: 1240K capture data for 22 individuals from the Kondopoga cemetery on Lake Onega, a churchyard used from the 12th to the 19th century. Combining relatedness estimates, autosomal and X-chromosomal identity-by-descent, uniparental lineages, burial position and archaeological dating, we place eight individuals in a single pedigree spanning five generations, anchored on a founding couple buried in the highest part of the cemetery. Identity-by-descent links almost all the remaining individuals to that couple, indicating that the sample is not a cross-section of a local population but a single kindred, which is the object of this study. Within it, the founding male’s paternal lineage recurs from the 12th–13th to the 16th century, mitochondrial lineages differ between individuals of the same generation, and autosomal ancestry shifts between generations rather than between archaeological phases. The ancestry of the founders is closest to populations from northwestern Rus’ and the eastern Baltic, while the Uralic-related ancestry entering the pedigree is best modelled using present-day Finnic groups of the region. The family was patrilineal, and its profile was reshaped by the women who married into it.

**Impact statement:** A five-generation pedigree from a Karelian churchyard shows a patrilineal kindred with Balto-Slavic-associated founders, whose genetic profile was remade, generation by generation, by women from the surrounding Finnic-speaking population.

## INTRODUCTION

Genome-wide data from burial grounds have made it possible to move from describing the ancestry of past populations to reconstructing the kinship systems of individual communities. Pedigrees from individual sites have revealed patterns of descent, residence and social inheritance, from Neolithic tombs and Bronze Age farmsteads to early medieval cemeteries. (Schroeder et al., 2019; Mittnik et al., 2019; Fowler et al., 2022; Rivollat et al., 2023; Gnecchi-Ruscone et al., 2024). Two recent methodological advances have made these analyses more precise. Relatedness estimators designed for pseudo-haploid, low-coverage data can identify first-, second- and third-degree relatives and distinguish parent–offspring from full-sibling pairs (Alaçamlı et al., 2024), while imputation-based detection of identity-by-descent segments can identify more distant relatives, from fourth to sixth degree, and trace connections across several generations (Ringbauer et al., 2024).

Combined with runs of homozygosity, which record parental relatedness and the size of the mating pool (Ringbauer et al., 2021), and with uniparental markers, these methods can convert a set of burials into a genealogy. Sharing on the X chromosome adds a further constraint, because a man inherits his X chromosome only from his mother and a father transmits a single, non-recombined X chromosome to every daughter; configurations of equal autosomal degree can therefore be distinguished according to whether the connection passes through a male or a female ancestor.

Most pedigrees of this kind come from sites used over a few generations, and reconstructions that span many centuries at one place remain uncommon. Long-lived churchyards are particularly informative in this respect, because continuity of the burial place can be compared directly with continuity of descent: a cemetery used by one lineage for centuries should show both a persistent paternal line and a recurrent influx of unrelated individuals through marriage, whereas a cemetery serving a changing community should not. A recent study of medieval Finland documented fine-scale genetic continuity together within such a setting (Nordfors et al., 2025), but for northern European Russia no comparable reconstruction has been available. No medieval individual from Karelia is represented in the available ancient DNA record at all: the only published ancient genomes from the region are Mesolithic, from Yuzhny Oleniy Ostrov (Fu et al., 2016; Mittnik et al., 2018; Posth et al., 2023), and the closest published samples of comparable date come from sites of medieval Rus’ in the Volga–Oka interfluve (Peltola et al., 2023) and in present-day Vologda Oblast (Andreeva et al., 2025).

The Kondopoga necropolis offers an opportunity of this kind. It was revealed during salvage excavations at the site of the Assumption Church on Kondopoga Bay, Lake Onega, after the wooden church standing there was deliberately burned down in an act of vandalism in 2018 (Reshetova and Nikitina, 2025). The excavations uncovered a necropolis used from the 12th to the 19th centuries. Its earliest graves, dated to the 12th–15th centuries, are the oldest medieval burials currently known in Olonets Karelia and precede the first written mention of Kondopoga in the cadastral books of the Obonezhskaya pyatina (Pokrovskii, 1930). Burial continued in the same plot over several centuries, alongside a succession of churches that stood at the Kondopoga churchyard from at least the 16th century (Suslova, 2013; Chernyakova et al., 1999). The graves are not randomly distributed: the earliest of them occupy the highest, central part of the hill, and one is a stone cist accompanied by gold and silver costume elements.

In its wider setting, the Lake Onega region was in the medieval period an area in which Baltic-Finnic-speaking populations, documented by a dense layer of Baltic-Finnic toponymy, and Slavic-speaking populations associated with the Novgorod lands were both present (Makarov, 1997; Spiridonov, 2001a, 2001b). The ancestry of individuals entering a local family through marriage is therefore an informative observation in its own right, independently of any claim about the regional population as a whole.

Here we reconstruct the genealogy of the community buried at Kondopoga. We first establish first- to third-degree relationships and anchor the pedigree on the earliest burials, then extend it with autosomal and X-chromosomal identity-by-descent to more distant relatives, and test its internal consistency against uniparental haplogroups, burial position and archaeological dating. We then ask three questions about the resulting kindred: whether the paternal line persisted, whether partners were drawn from within or outside the local relatedness pool, and whether the genetic profile of the family changed along the pedigree. We use population-genetic analyses, including principal component analysis, ADMIXTURE, f3 and f4 statistics, and qpAdm, to characterise the ancestry of individuals in the pedigree and examine how it changed across generations. Our focus is on ancestry within this family, rather than on regional population history. We find that almost the entire sampled cemetery belonged to a single family descended from one couple. Its paternal lineage persisted over several centuries, while successive generations introduced new maternal lineages and showed changes in autosomal ancestry.

## RESULTS

We screened 45 libraries from 32 individuals, including adults and children, from the Kondopoga burial site in the Republic of Karelia, Russia (Figure 1), for ancient DNA preservation. Twenty-three libraries from 23 individuals showed >1% endogenous DNA and characteristic deamination patterns and were enriched using a 1.24 million SNP capture panel. Post-capture data from 22 individuals (one individual was excluded because of high contamination; Supplementary Materials, Table S1, Supplementary Figures, Figure S1) contained a mean of 646,551 SNPs per individual (range: 254,775–963,996), providing sufficient coverage for IBD and hapROH analyses in most samples.

**Figure 1.**
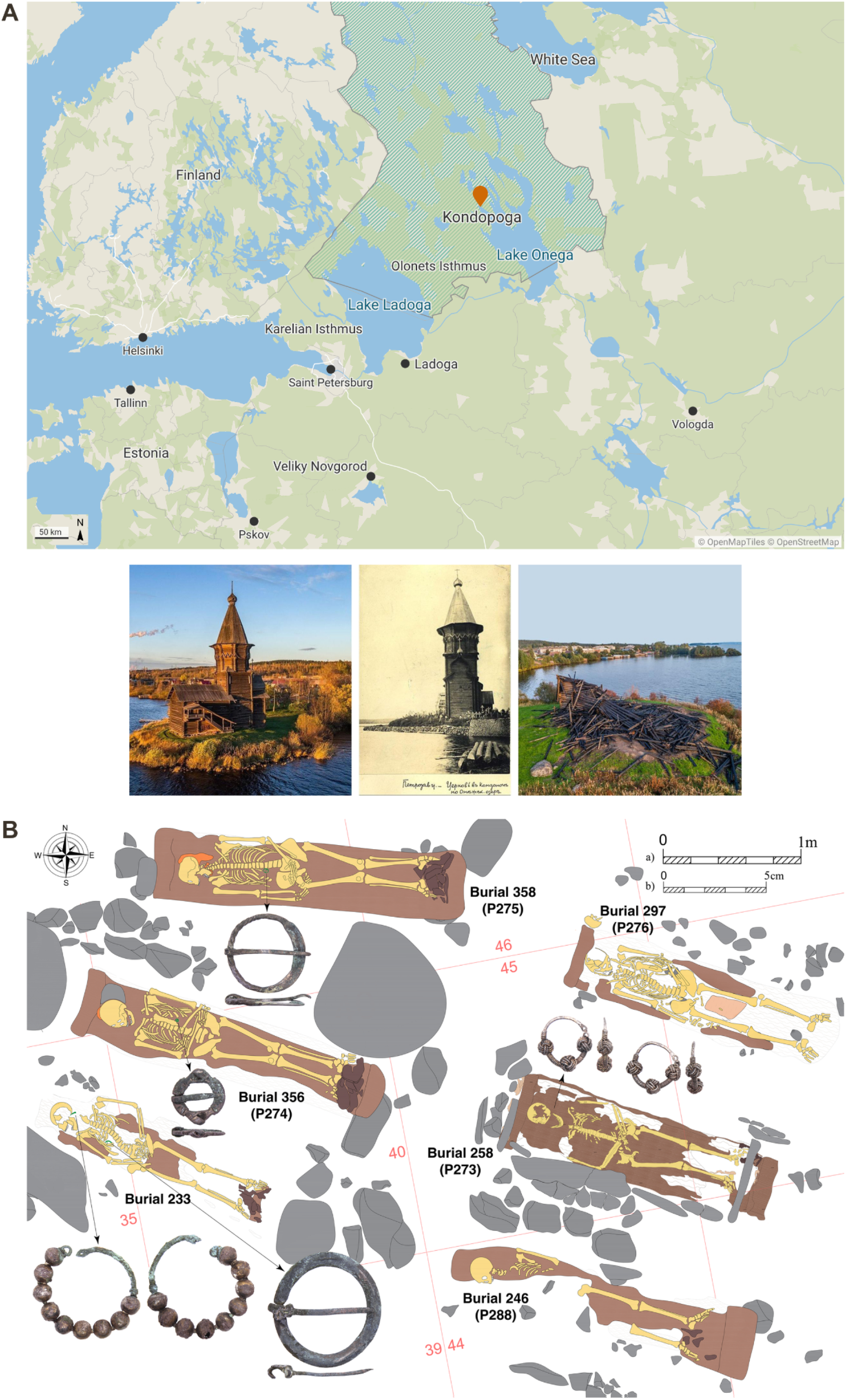
The Kondopoga necropolis and its setting. **(A)** Location of Kondopoga on Kondopoga Bay, Lake Onega (marker); the hatched area is the modern Republic of Karelia. Base map © OpenMapTiles, © OpenStreetMap contributors. Below, the Assumption Church of Kondopoga, at the site of which the cemetery was excavated: the church before the fire (left), on an early twentieth-century postcard (centre), and the remains after it was burned down in 2018 (right). The photographs of the church are provided by the Institute of Archaeology, Russian Academy of Sciences (IA RAS). **(B)** Plan of the central part of the burial ground, showing the graves of the founding couple P273♀ (burial 258) and P276♂ (burial 297), their sons P274♂ (burial 356) and P288♂ (burial 246) and their grandson P275♂ (burial 358), together with burial 233 and the associated grave goods. Scale bars: a) 1 m, b) 5 cm.

### A founding couple and a five-generation pedigree

Genetic sex was in full agreement with the sex assigned through archaeological methods, giving 10 females and 12 males. All 22 individuals passed the coverage threshold required for identity-by-descent analysis and 19 had the ≥400,000 SNPs required for the analysis of runs of homozygosity, so the great majority of the burial community could be placed in the genealogical reconstruction that follows. Among the 22 individuals, 18 carried subclades of mitochondrial haplogroup H, two haplogroup T2b, and one each J1b1a1 and V7a1; among the 12 males, nine carried lineages of R1a - seven R-L1029 and its subclades R-YP417 and R-YP418, one the upstream R-PF6155 and one the separate branch R-YP5520 - while two carried I-Y3120 and one N-Y16503 (Supplementary Materials, Table S1). The single N1a lineage belongs to one of the earliest individuals at the site and is not represented in any later generation. The distribution of these lineages among other ancient and present-day populations is surveyed in Supplementary Note 1.

We estimated biological relatedness for all 231 pairs with READv2 (Supplementary Materials, Table S3). Twenty pairs were assigned a relationship: seven of the first degree, six of the second and seven of the third. The first-degree assignments are internally consistent with the uniparental data. The mother–son relationships inferred for P273♀ with P274♂ and P288♂ are consistent with their shared mitochondrial haplogroup H, and the father–son relationship inferred for P276♂ with P288♂ is consistent with their shared Y-chromosomal haplogroup R-YP418*. P273♀ and P276♂ are therefore the parents of at least two sampled sons, and P275♂ is the son of P274♂ and thus their grandson. The direction of descent was fixed independently of the genetic data by the archaeological dating of the graves and by the age at death estimated for each individual.

X-chromosomal identity-by-descent (Supplementary Materials, Table S16) was used alongside the autosomal estimates to distinguish between genealogical configurations of equal autosomal degree. The pedigree satisfies the expectations of X-chromosomal transmission at every link used in the reconstruction. Every male–male relative pair shares no X chromosome at all - P276♂–P288♂, P274♂–P276♂, P274♂–P275♂ and P298♂–P300♂ all share 0 cM despite 2,528–3,216 cM of autosomal sharing - as expected because a man transmits his Y chromosome, not his X, to his sons. Mother–son pairs share the most X chromosome at the site (P273♀–P274♂, 176 cM; P273♀–P288♂, 158 cM), the brothers P274♂ and P288♂ share 62 cM through their mother, and P273♀ and her grandson P275♂, whose connection passes through her son, share none.

Two further individuals can be placed in the pedigree, and in each case X-chromosomal sharing also resolves the sex of the unsampled individual through whom the connection passes. P285♀ is assigned a second-degree relationship to P275♂, a third-degree relationship to P274♂ and fourth-degree levels of sharing with P276♂ and P288♂ (329 cM and 316 cM), the combination expected of a grandchild of P275♂; she shares no X chromosome with P275♂ despite 1,260 cM of autosomal sharing, so her descent passes through a son of P275♂ rather than a daughter, and her mitochondrial haplogroup H28a accordingly differs from his H1n4; the 14.3 cM she shares with P273♀ and with P276♂ is at the median level of X-chromosomal sharing among all pairs at the site (15.4 cM) and does not constrain her position. P282♂ is assigned a second-degree relationship to P288♂ and third-degree relationships to P276♂ and P274♂, the combination expected of a grandson of P288♂; he shares no X chromosome with P273♀, P276♂, P274♂ or P288♂, so his own X chromosome derives entirely from his mother, and his descent from P288♂ therefore passes through a son. Taken together with the third-degree relationships, 11 of the 22 individuals - P273♀, P274♂, P275♂, P276♂, P282♂, P283♀, P285♀, P288♂, P293♀, P295♀ and P297♂ - belong to one extended family. Eight of them can be given a position in a pedigree spanning five generations; the positions of P283♀, P293♀ and P297♂ remain unresolved (Figure 2). P273♀ and P276♂ can accordingly be treated as the founding couple of the genetically reconstructed pedigree.

**Figure 2.**
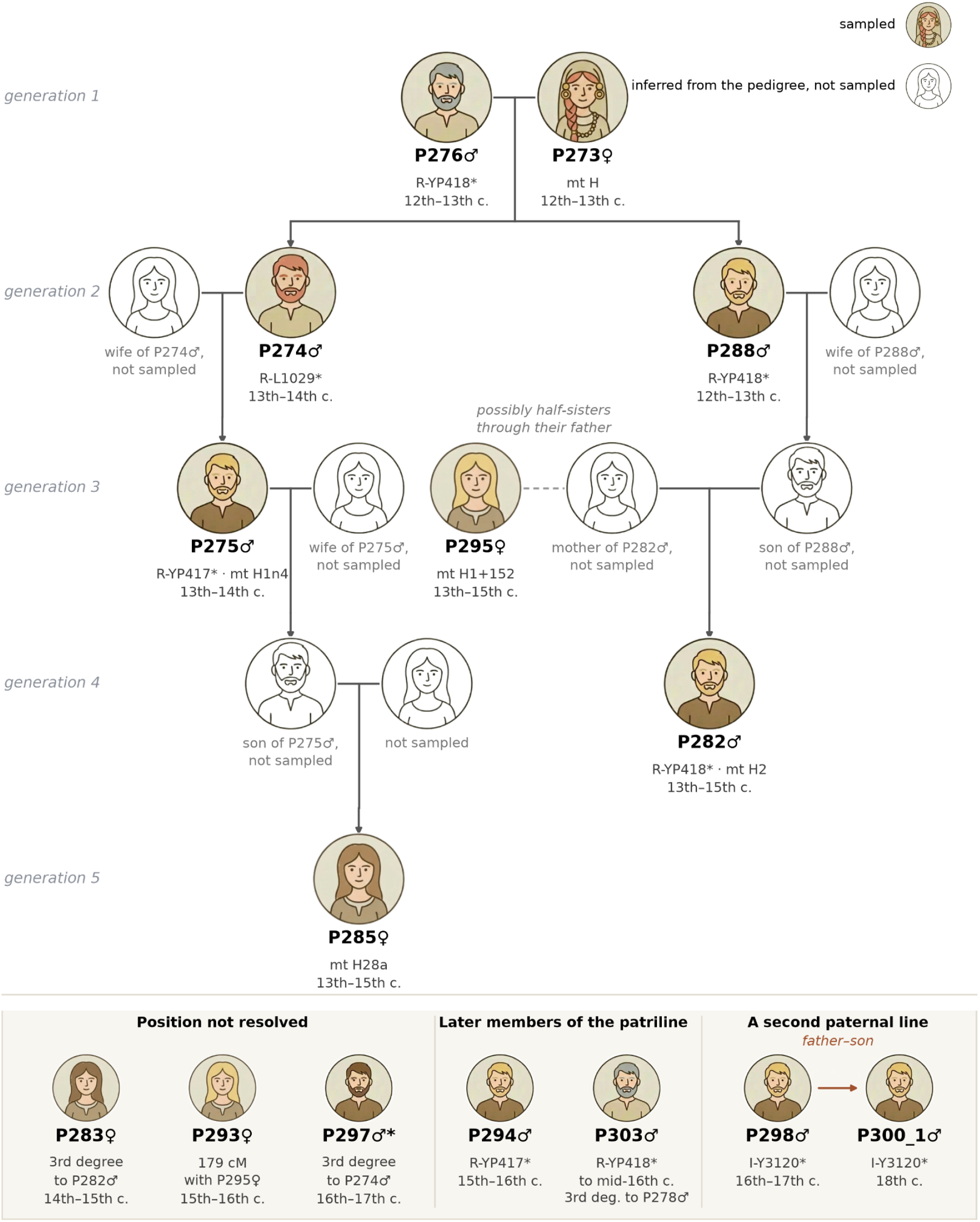
Pedigree of the Kondopoga kindred. Genealogy reconstructed from READv2 relatedness estimates, autosomal and X-chromosomal identity-by-descent sharing, uniparental haplogroups, age at death and the archaeological dating of the burials. Individuals are arranged by generation; shaded portraits are sampled individuals, open portraits individuals required by the reconstruction but not sampled. Y-chromosomal and mitochondrial haplogroups and the dating of the burial are given below each sampled individual. P285♀ is placed as a granddaughter of P275♂ and P282♂ as a grandson of P288♂, in both cases through an unsampled son, because neither shares an X chromosome with the paternal line; P295♀ and the unsampled mother of P282♂ are drawn as possible half-sisters through their father (dashed line), the configuration consistent with their X-chromosomal sharing and their differing mitochondrial lineages. The band at the foot of the figure shows individuals related to the kindred but not placed within it: three whose position is not resolved, two later members of the founding patriline, and the second paternal line formed by the father–son pair P298♂ and P300_1♂. Hair colour of the sampled individuals follows the HIrisPlex predictions in Supplementary Table S17 and grey hair the age at death recorded in the archaeological documentation; portraits are schematic and are not reconstructions of appearance. The third-degree relationship of P297♂ to P274♂ (813 cM; longest shared segment 99 cM) places him three generations after P274♂, whereas his burial is dated to the 16th–17th centuries; the two assignments cannot both be correct and the discrepancy is unresolved. P279♀, P286♂, P290♀, P299♀, P301♀ and P302_1♀ share no more than 87 cM with any member of the pedigree, consistent with background relatedness in a small mating pool (effective population size 2,875; 95% CI 2,092–4,063).

The pedigree is also consistent with the Y-chromosomal data. P274♂ was assigned to R-L1029* and P275♂ to R-YP417*, both ancestral to the R-YP418* lineage carried by P276♂ and P288♂; because P274♂ and P275♂ lacked the coverage required for assignment to a more derived subclade, the male line P276♂ → P274♂ → P275♂ is consistent with continuity of R-YP418 across three successive generations rather than with the entry of a different paternal lineage.

### Identity-by-descent extends the genealogy to almost the whole cemetery

Relatedness beyond the third degree cannot be resolved with pseudo-haploid relatedness estimators, so we called identity-by-descent segments with ancIBD on imputed genotypes for all 22 individuals. The results reproduced every first- to third-degree relationship inferred with READv2 (Supplementary Materials, Table S15; Figure 3A) and additionally resolved fourth-, fifth- and sixth-degree pairs (Figure 3B), which extends the reconstruction by a further two to three generations in each direction. Sharing is pervasive: 149 of the 231 pairs share at least one segment longer than 12 cM (Figure 3C).

**Figure 3.**
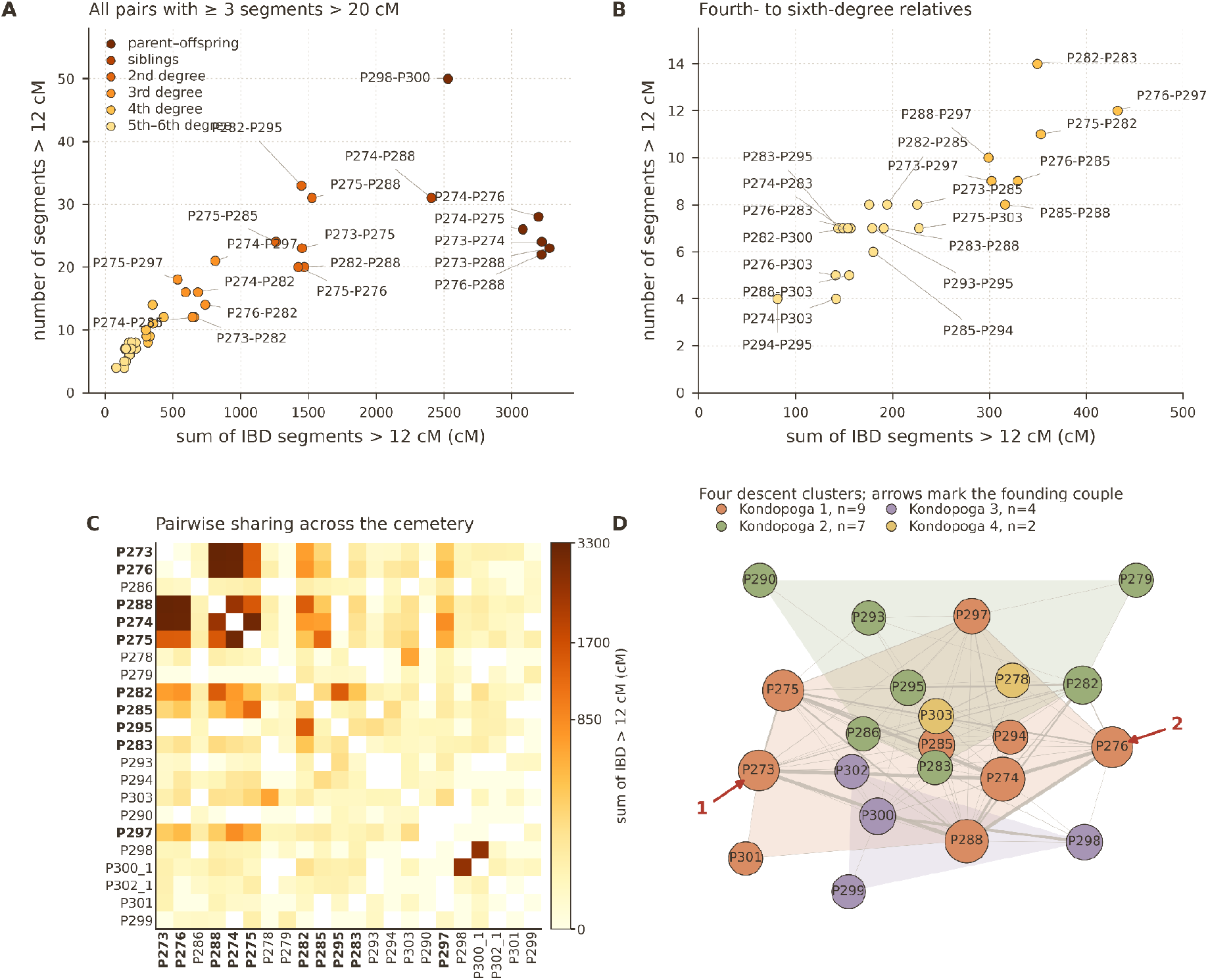
Identity-by-descent (IBD) sharing among Kondopoga individuals. **(A)** Number and total length of IBD segments longer than 12 cM for every pair sharing at least three segments longer than 20 cM. Colours give the degree of relatedness inferred from the IBD data; first-degree pairs are separated into parent–offspring and siblings using READv2 and X-chromosomal sharing. Labels are given for pairs involving a member of the founding family and for P298♂–P300♂. **(B)** The same quantities for the fourth- to sixth-degree pairs, on an expanded scale. **(C)** Total length of IBD segments longer than 12 cM for all 231 pairs, ordered by archaeological date; individuals placed in the pedigree are in bold. **(D)** Clustered IBD network of the Kondopoga individuals. Nodes represent samples and edges pairwise sharing above 20 cM, with edge width proportional to the total length shared. Colours denote the four clusters identified by the Leiden algorithm (resolution parameter = 1) and convex hulls their boundaries; the layout was generated with the ForceAtlas2 algorithm. Red arrows indicate the earliest Kondopoga individuals: P273♀ (1) and P276♂ (2). The corresponding network including 19 individuals from northern Rus’ is given in Supplementary Figures, Figure S2.

The reach of the founding couple within the cemetery is the clearest expression of this structure. P273♀ and P276♂ share segments longer than 20 cM with 15 of the 20 other sampled individuals, and 19 of the 20 share at least one segment longer than 8 cM with one of them. Clustering the network of pairwise sharing above 20 cM with the Leiden algorithm partitions the Kondopoga individuals into four descent clusters, of which the founding couple and their immediate descendants form the core of the largest; the remaining clusters are connected to it through fourth- to sixth-degree links rather than forming separate components (Figure 3D). The sampled burial community is therefore not a cross-section of a local population but a single kindred, traceable over the whole period during which the churchyard was in use, together with a small number of individuals attached to it by more distant ties.

The in-marrying side of the family is itself represented at the cemetery, and X-chromosomal sharing distinguishes it from the line of descent. P295♀ shares no identity-by-descent segment with either founder, yet shares 1,448 cM in 33 segments longer than 12 cM with P282♂ and is assigned a second-degree relationship to him. The pair also shares 146 cM of X-chromosomal identity-by-descent, the highest value recorded for a male–female pair at the site outside the mother–son pairs, which places the connection on P282♂’s maternal side. Her mitochondrial haplogroup H1+152 differs from his H2, which excludes a shared maternal line and therefore excludes her being his maternal grandmother or his mother’s full sister. The configuration that satisfies all three observations is that P295♀ and the mother of P282♂ were half-sisters through their father, since a father transmits a single, non-recombined X chromosome to every daughter while different mothers give different mitochondrial lineages. P295♀ is thus a relative of a woman who married into the lineage rather than a descendant of the founders, and she carries the highest burden of background relatedness in the dataset (37.6 cM in runs of homozygosity longer than 4 cM, against no segment above 2.7 cM in P273♀), consistent with local origin. P293♀ and P283♀, who share 179 cM and 144 cM with her respectively, appear to belong to the same side of the family.

The four clusters differ in composition in a way that follows the structure of the kindred. The largest (n = 9) is the core of the family: the founding couple, their sons P274♂ and P288♂, P275♂, P285♀ and the later P294♂ and P297♂, every male of whom carries a Y-chromosomal haplogroup of the R1a1a1b1a1 branch; its members share on average 846 cM with one another and only 78 cM with the rest of the cemetery. P301♀ of the eighteenth–nineteenth centuries also falls here, but at a mean of only 19 cM. The second cluster (n = 7) consists mainly of women, and it is here that P295♀, P293♀ and P283♀ fall, together with P279♀ and P290♀; each of the five carries a different mitochondrial haplogroup (H1+152, H1+16278, J1b1a1+146, V7a1 and H1a2), and they appear with P282♂ of the founding patriline and P286♂, the only male in the cemetery carrying N-Y16503*, a lineage of Siberian origin that is absent from every later male. Mean sharing within this cluster is 146 cM, the level of fourth- to sixth-degree relatives, which is what is expected of women who entered the kindred from different families rather than descending from a common ancestor within it. The third cluster (n = 4) is the latest and the most weakly attached to the rest of the cemetery (mean 27 cM): it holds the second paternal line, P298♂ and P300♂ (I-Y3120*), and two women, P302_1♀ and P299♀, all dated between the sixteenth–seventeenth centuries and the nineteenth. The fourth cluster is the pair P278♂ and P303♂, third-degree relatives sharing 594 cM whose paternal lineages nevertheless differ: P303♂ carries the R-YP418* of the founding patriline, whereas P278♂ belongs to a separate R1a branch, R-YP5520*.

One pair stands apart from this structure. P298♂ and P300♂ share 50 identity-by-descent segments longer than 12 cM, the highest number in the dataset, and their inferred father–son relationship is consistent with their shared Y-chromosomal haplogroup I-Y3120*; the direction of the relationship was assigned from the archaeological dating, with P298♂ as the father. Neither individual has a first-, second- or third-degree relative elsewhere in the dataset, and in the identity-by-descent network they fall into a separate late cluster. They therefore represent a second, shorter paternal line using the same burial ground rather than a branch of the main kindred.

Two individuals extend the paternal lineage into the sixteenth century. P303♂, whose burial is dated to no later than the middle of the sixteenth century, carries R-YP418*, the same subclade as the founding male P276♂, his son P288♂ and his fourth-generation descendant P282♂, and P294♂ (fifteenth–sixteenth centuries) carries the ancestral branch R-YP417*. Their autosomal sharing with the founding generations has decayed to fifth- and sixth-degree levels: P303♂ shares 227 cM with P275♂, 155 cM with P288♂ and 141 cM with each of P274♂ and P276♂, and P294♂ shares 136 cM with P275♂ and 116 cM with P274♂. The closest relationship of P303♂ is a third-degree one to P278♂ (594 cM, longest shared segment 112 cM), whose own sharing with the founding generations does not exceed 68 cM. These individuals therefore belong to a collateral branch of the same patrilineage: the Y-chromosomal lineage of the founding male is still present some three centuries after his burial, at a point where autosomal relatedness to him has diluted to fifth- and sixth-degree levels. An undiminished paternal haplogroup alongside decaying autosomal sharing is the signature expected of a patrilineage that continued to use one burial ground over many generations, and it is the strongest evidence we have for continuity of the male line (Figure 4A).

**Figure 4.**
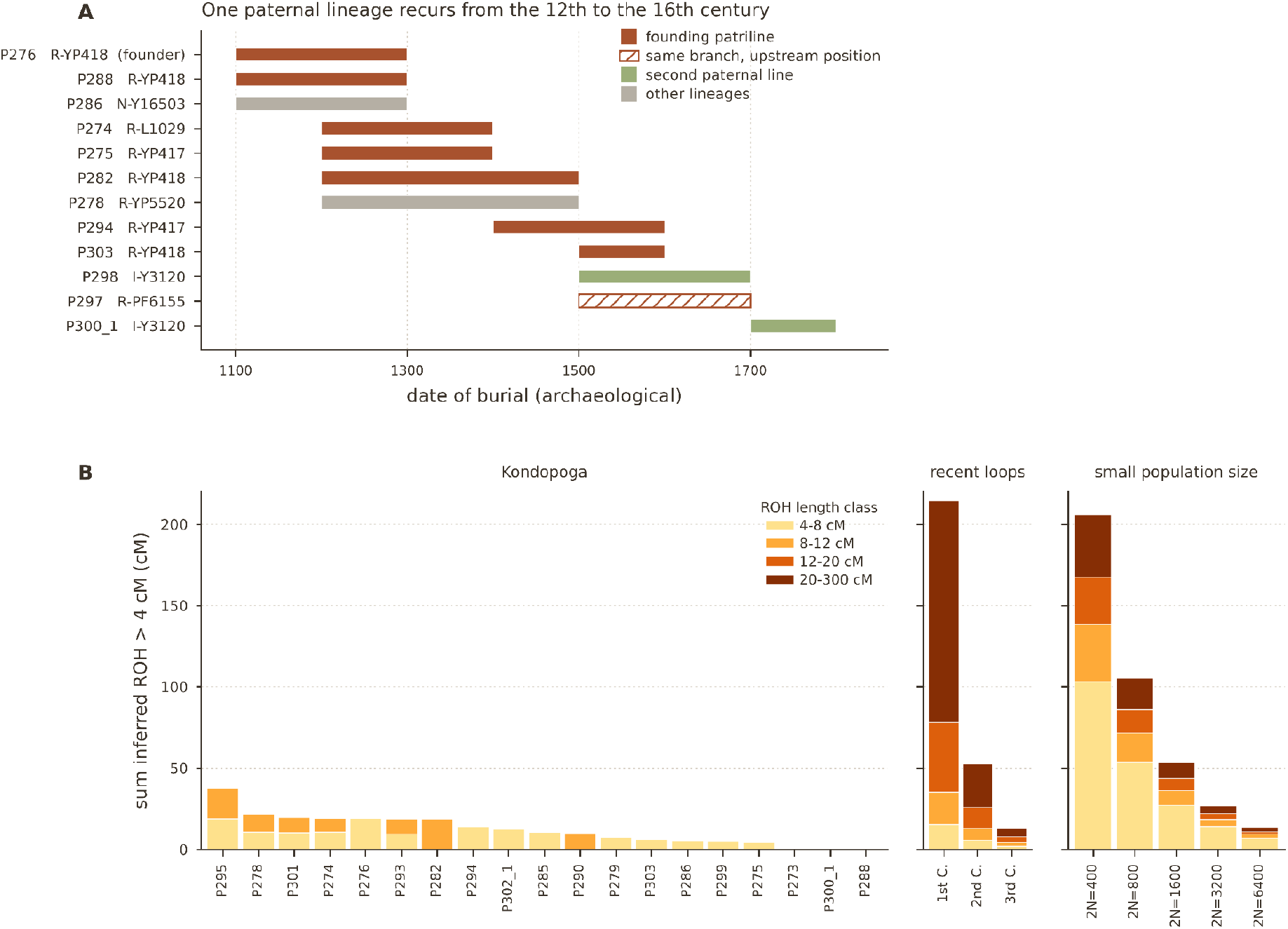
Paternal lineages and background relatedness. **(A)** Y-chromosomal haplogroups of the twelve males, plotted against the archaeological date of their burials; each bar spans the dating interval of the grave. Colour separates the founding patriline from the second paternal line and from unrelated lineages. The hatched bar marks P297♂, whose haplogroup R-PF6155* lies upstream of the founding patriline within R1a1a1b1a1 and is therefore compatible with membership of it without establishing it. **(B)** Runs of homozygosity inferred with hapROH for the 19 individuals with sufficient coverage, given as the summed length of segments longer than 4 cM and split into length classes. The two blocks on the right give the profiles expected for the offspring of first, second and third cousins and for constant population sizes of 2N = 400 to 6,400, computed with the same package.

One individual cannot be reconciled with the chronology. P297♂ (burial 323) is assigned third-degree relationships to P274♂ (813 cM in 21 segments longer than 12 cM, the longest 99 cM) and to P275♂ (534 cM), together with fourth- to fifth-degree levels of sharing with P276♂ (432 cM), P273♀ (302 cM) and P288♂ (299 cM). Taken together these values match the expectation for a great-grandchild of P274♂ descending through a child other than P275♂ (870, 430, 430, 430 and 200 cM respectively), which would place him in the late fourteenth or the fifteenth century, whereas burial 323 is dated to the sixteenth–seventeenth centuries. The genetic signal is unlikely to be an artefact of accumulated background relatedness, which produces numerous short segments rather than segments approaching 100 cM, and the contamination estimates for this library are low (nuclear 0.34–0.39%, mitochondrial 0.02; Supplementary Materials, Table S2), although the nuclear estimate rests on only 1,851 informative SNPs and this individual has the lowest mitochondrial haplogroup assignment quality in the dataset (0.75). Burial 323 also has the least constrained archaeological date among the graves discussed here: it was identified in the cemetery layer in a shallow pit with amorphous outlines and bears no stratigraphic relation to another burial or to a dated structure, unlike burials 182, 208 and 228, each of which is tied stratigraphically to an adjacent grave, and its date rests on a single costume item, a copper-alloy finger ring with a hollow hexagonal bezel. The grave is also one of only two in the cemetery oriented with the head to the south-south-east rather than following the alignment of the church. We report the discrepancy rather than resolve it, and treat P297♂ as a member of the kindred without a position in the pedigree. His Y-chromosomal haplogroup does not contradict membership of the founding patriline: R-PF6155* sits at R1a1a1b1a1, upstream of the R-L1029*, R-YP417* and R-YP418* carried by the men of that line, so the call is consistent with it but too shallow to be diagnostic. The correspondence between library identifiers and burial numbers is given in Supplementary Materials, Table S1.

### No close-kin unions and a small mating pool

A kindred that used one burial ground for several centuries might be expected to show the effects of marriage within it, and runs of homozygosity test this directly. We detected short runs of homozygosity (ROHs) in nearly all 19 individuals analysed, but none of the individuals carried long ROHs (>20 cM) that would indicate close parental relatedness within the last few generations (Figure 4B; Supplementary Materials, Table S4). This pattern is consistent with distant background relatedness in a population with a relatively small effective population size. The effective population size was estimated at 2,875 individuals (95% confidence interval [CI]: 2,092–4,063). Because the estimate integrates genetic information across several centuries, it should not be interpreted as the census size of the Kondopoga settlement, but rather as an estimate of the effective population contributing to the sampled individuals.

Notably, the earliest sample, P273♀, showed no ROH segments >4 cM, which may indicate lower levels of distant parental relatedness than observed in the other Kondopoga individuals. This pattern is also consistent with the possibility that P273♀ originated from a larger population or settlement where distant parental relatedness was less common. Additionally, no ROH segments longer than 4 cM were detected in P288♂, the son of P273♀, or in P300_1♂, one of the males carrying the I-Y3120* haplogroup. The second individual carrying this haplogroup was not analysed by this method because he had fewer than 400,000 SNPs.

### Burial position follows the pedigree

Burials P273♀ and P276♂ have been identified by archaeologists as the core of the burial ground: they are among the earliest graves, dated to the 12th–13th centuries, located in the highest central part of the hill, and may have held particular symbolic significance. Burial P273♀ was interred in a stone cist constructed from vertically arranged flat stone blocks and was accompanied by gold and silver costume elements and ornaments, which may indicate a high social status (Archaeological Supplementary, burials 258 and 297, Figure 1B). These two burials (P273♀ and P276♂), together with their son P288♂, also dated to the late 12th–13th centuries, form a spatially and genealogically coherent group. The burial of another son, P274♂, and that of a grandson, P275♂, both dated to the 13th–14th centuries, are also located in close proximity (Figure 1B). The spatial arrangement of these graves therefore follows the reconstructed genealogy, with the founding couple and their closest descendants occupying the centre of the cemetery.

### Ancestry of the kindred and its change across generations

Having established how the individuals buried at Kondopoga were related to one another, we turn to their ancestry and to how it changed from one generation of the kindred to the next. To compare Kondopoga samples to present-day populations, we performed a Eurasian-wide principal component analysis (PCA) with 79 present-day populations (Figure 5, Supplementary Materials, Table S5). The positioning of the samples shows variation within the group, but overall, all samples fall within the range of present-day North Russian and Finno-Karelian populations, particularly Karelians and Veps. P273♀ and her sons P274♂ and P288♂ are positioned closer to present-day Estonians and Lithuanians than the other Kondopoga individuals, indicating a distinct position within the Kondopoga group.

**Figure 5.**
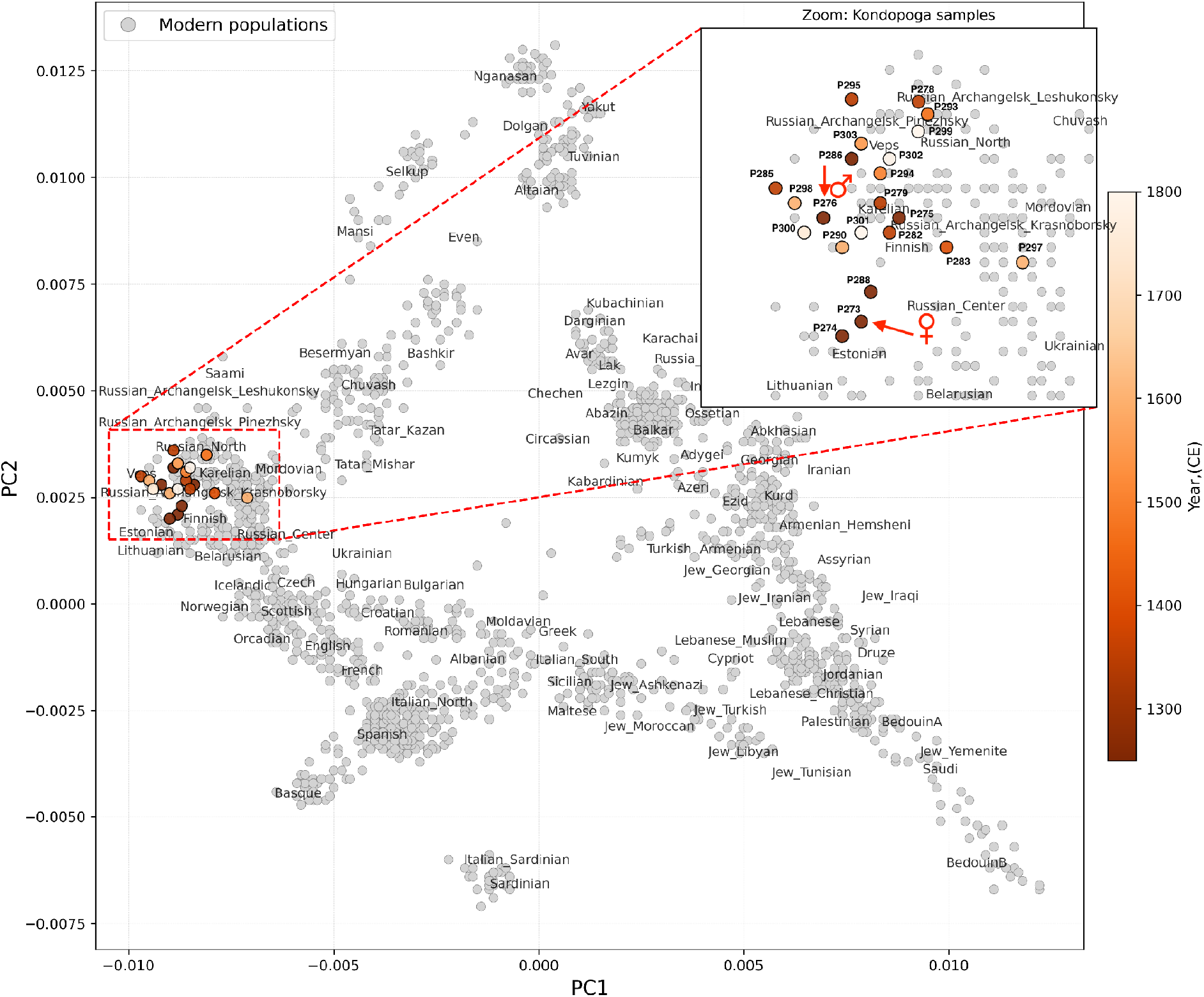
Principal component analysis of the Kondopoga individuals against present-day Eurasian populations. Grey points are individuals from 79 present-day populations (Supplementary Materials, Table S5); coloured circles are the 22 Kondopoga individuals, with the colour scale giving the archaeological date of the burial. Red arrows mark the founding couple, P273♀ and P276♂. The inset enlarges the region of the plot occupied by the Kondopoga individuals, outlined by the dashed red rectangle in the main panel, and labels each of them.

Projection of the Kondopoga individuals together with ancient groups of comparable date and region reproduces this division (Supplementary Figures, Figure S3). The earliest individuals - P273♀ and her sons P274♂ and P288♂ - fall among Iron Age and medieval groups of the eastern Baltic and of the Slavic-period west, whereas most of the later individuals plot with medieval individuals from northwestern Rus’ and with groups carrying Finnic-associated ancestry. A regional principal component analysis against 123 individuals from 15 Baltic- and Finnic-related populations of northwestern Russia (Agdzhoyan et al., 2024) shows the same pattern: the founding family plots with present-day Russian populations of the Pskov, Yaroslavl and Novgorod regions, the majority of the Kondopoga individuals fall within the Karelo-Finnic cluster, and P276♂ occupies an intermediate position between the two (Supplementary Figures, Figure S4). The named comparison groups and the position of each individual are given in Supplementary Note 2 (Supplementary Materials, Tables S5 and S6).

The ancestry components themselves show the same contrast. ADMIXTURE analysis at K = 6 (Figure 6A) revealed three major components in the Kondopoga population corresponding to Early European Farmer (EEF), Western Hunter-Gatherer (WHG), and Ancient North Eurasian (ANE)-related variation. These components are widely distributed among present-day European populations (Lazaridis et al., 2014). In addition, a fourth component with a higher frequency in present-day Asian populations is present in the Kondopoga individuals. We interpret this component as reflecting Uralic-related ancestry, which is also present in present-day Finnic populations (Zeng et al., 2025).

**Figure 6.**
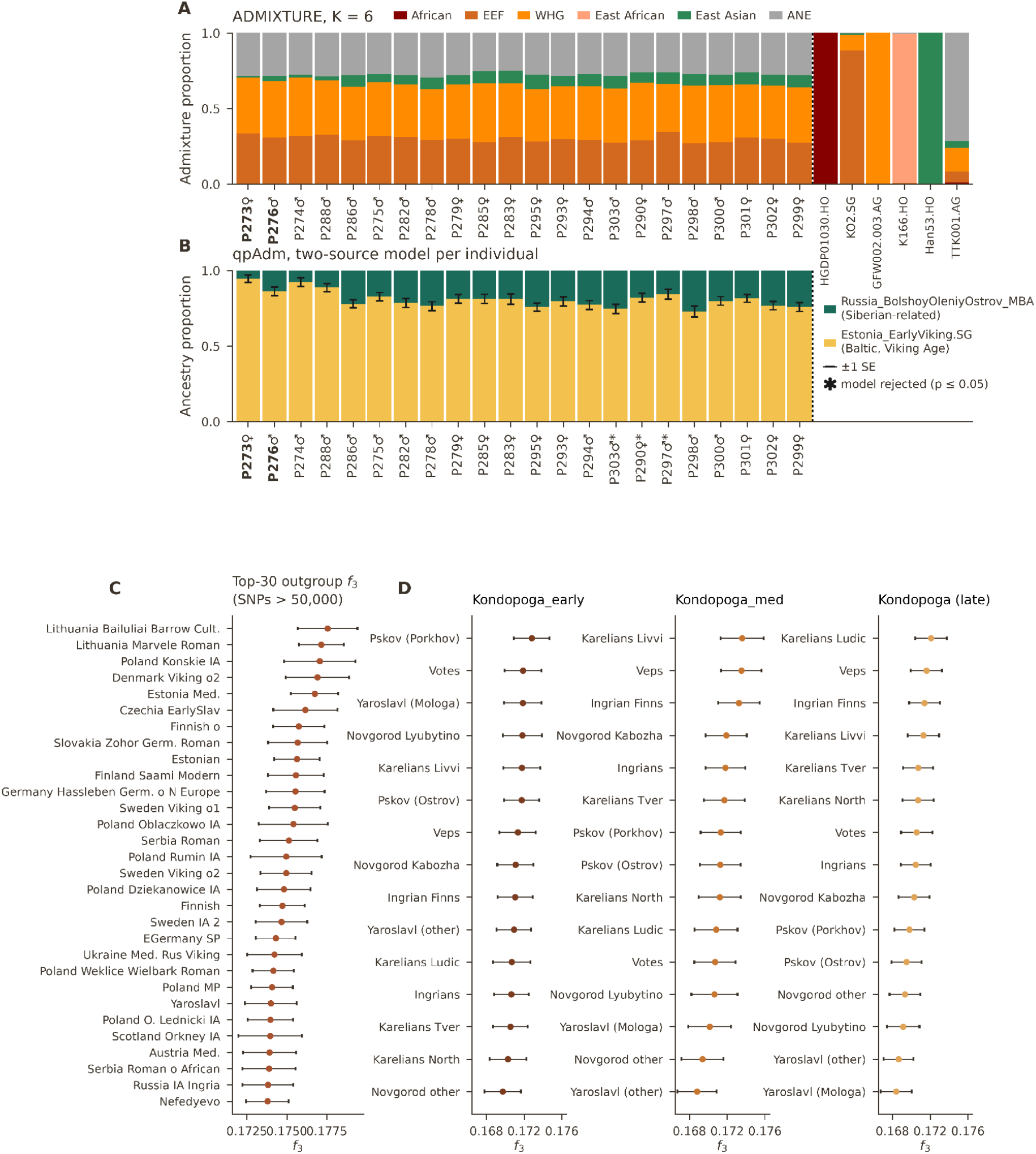
Ancestry of the Kondopoga individuals and its change across generations. **(A)** ADMIXTURE at K = 6 for the 22 Kondopoga individuals, with six reference individuals to the right of the dotted line anchoring the components. Individuals are ordered by the archaeological date of the burial; the symbol after each sample name gives the sex, and the founding couple P273♀ and P276♂ is set in bold. **(B)** qpAdm two-source models fitted to each individual separately, with Estonia_EarlyViking.SG and Russia_BolshoyOleniyOstrov_MBA as sources and ten right populations (Methods). Bars give the estimated proportions, the black whisker is ±1 SE on the boundary between the two components, and an asterisk marks the three individuals whose model is rejected (p ≤ 0.05). Individual order and bold type are as in **(A)**; full results for all four source pairs tested are given in Supplementary Materials, Table S14. **(C)** The thirty highest outgroup-f3 values f3(Kondopoga, X; Yoruba.HO) among tests covering more than 50,000 SNPs. **(D)** Outgroup-f3 computed separately for the three temporal groups, fifteen populations each, ordered by f3 within each group. f4 statistics are given in Supplementary Figures, Figure S9.

The Uralic-related component was less pronounced in the earliest pair (P273♀ and P276♂) and their sons (P274♂ and P288♂) than in most of the other Kondopoga individuals. In P275♂, the grandson of the founding couple, the proportion of this component becomes comparable to that observed in the other Kondopoga individuals. This pattern may reflect admixture between the early Kondopoga population and populations carrying a higher proportion of Uralic-related ancestry. The proportion of this component increased over the subsequent 3–4 generations and subsequently remained relatively stable (Figure 6A).

The comparison between P274♂ and his son P275♂ is consistent with a substantial contribution of Uralic-related ancestry from the unsampled mother of P275♂. P282♂, a grandson of P288♂ and representing the fourth generation in the reconstructed pedigree, carries an even higher proportion of Uralic-related ancestry than P275♂. Together, these results are consistent with increasing Uralic-related ancestry among descendants of the founding couple and may reflect repeated admixture with populations associated with Finnic-speaking communities of the region and carrying a higher proportion of Uralic-related ancestry.

The ADMIXTURE profile of the founding family (P273♀, P276♂, P274♂ and P288♂) matches those of individuals from northwestern Rus’, of a subset of medieval Estonia and Iron Age Ingria, and of present-day Estonians and Russians outside Vologda Oblast, whereas the remaining Kondopoga individuals carry proportions of the Uralic-related component comparable to present-day Karelians, Veps and Russians from Vologda and Arkhangelsk Oblasts. The proportion shows no decrease after the 13th century, including among the latest individuals in the dataset, which does not indicate a substantial additional contribution from populations of Central and northwestern Rus’ during this period (Supplementary Note 2; Supplementary Figures, Figures S5 and S6).

### Genetic affinities to other populations

The analyses above describe the cemetery from within; we next place its occupants among neighbouring, earlier and present-day populations. We computed outgroup-f3 statistics in the form f3(Kondopoga, Test; Yoruba.HO), where Test represented 643 ancient and present-day Eurasian populations dating to 2000 BP or later. The full results are presented in Supplementary Materials, Table S9, and the thirty populations with the highest f3 values among tests covering more than 50,000 SNPs are shown in Figure 6C. The highest f3 value among these tests was obtained for Lithuania_Bailuliai_BarrowCulture.SG, 0.1776, indicating substantial shared genetic drift between Kondopoga and ancient populations from the eastern Baltic region. Other highly ranked populations included Lithuania_Marvele_Roman.SG, Poland_Konskie_IA.SG, Estonia_Medieval.SG, Czechia_EarlySlav.AG, and several populations from Finland and Scandinavia. The thirty highest-ranked populations also included the medieval Nefedyevo group from the Vologda region, as well as EGermany_SP and Russia_IA_Ingria. Overall, the f3 results indicate strong shared genetic drift between Kondopoga and populations from the eastern Baltic and Central and Eastern Europe, consistent with the affinities observed in the PCA.

However, as the Kondopoga individuals span several centuries and potentially multiple generations, with possible admixture occurring during the settlement’s growth and development, we subdivided them into three temporal groups for further analysis: Kondopoga_early (the founding individuals: P273♀, P276♂, P274♂, P288♂), Kondopoga_med (their descendants from three subsequent generations: P275♂, P282♂, P285♀), and Kondopoga (the remaining samples with more distant or no direct kinship to the founding couple). We then performed outgroup-f3 statistics separately for the three groups in the form f3(Kondopoga_pop, Test; Yoruba.HO), where Test populations comprised modern Balto-Finnic groups from Russia (Agdzhoyan et al., 2024; Figure 6D; Supplementary Materials, Table S10). The results were fully consistent with the PCA findings. Kondopoga_early showed the highest levels of shared genetic drift with populations from Porkhov (Pskov region), Votes, and Yaroslavl, reflecting affinities to Baltic- and Slavic-related groups. In contrast, both later groups show their highest values with Finnic populations: Livvi Karelians, Veps and Ingrian Finns for Kondopoga_med, and Ludic Karelians, Veps and Ingrian Finns for the remaining individuals. Veps and Ingrian Finns are among the three highest-ranked populations in both groups, whereas Ludic Karelians rank first only in the latest group and tenth of fifteen in Kondopoga_med.

To detect individual-level differences, we also performed outgroup-f3 analyses for each sample separately, using Iron Age and Medieval test populations from relevant geographic regions, as well as modern populations from the HO dataset. For the first pair, P273 and P276, the highest f3 values were observed with Slavic- and Baltic-associated populations, with Medieval Poland (1000–1200 CE) ranking first. P273 also showed genetic affinity to Medieval Rus samples from Yaroslavl. Similarly, P274 and P288, the sons of P273 and P276, and P275, their grandson, did not show high affinity to Finnic populations or to North Medieval Rus groups (Nefedyevo, Minino, Nikolskoye and Voezero). In contrast, such affinities were observed in the majority of later samples. The results of the individual-level outgroup-f3 analyses with ancient and modern populations are presented in the Supplementary Figures, Figures S7 and S8 and Supplementary Materials, Table S11 and Table S12.

To assess changes in Finnic-related ancestry over time in the Kondopoga population, we performed f4 statistics in the form f4(Yoruba.HO, Test; Kondopoga_early, Kondopoga) (Supplementary Figures, Figure S9; Supplementary Materials, Table S13). We observed significantly positive values for populations carrying Eastern Eurasian and Siberian ancestry, including Nganasan (Z = 5.315), Han (Z = 5.091), Yakut (Z = 4.938), and Saami (Z = 3.108). In contrast, no significant signal was detected for Finnish, Karelian, Veps, or Russian populations (|Z| < 3). These results indicate increased affinity of the later Kondopoga individuals to Eastern Eurasian-related populations relative to the early group. Given the geographic distance from direct Eastern Eurasian sources, particularly East Asian populations such as Han, this signal is unlikely to reflect direct gene flow from these populations and may instead reflect indirect gene flow mediated by populations with mixed western and eastern Eurasian ancestry. The elevated f3 affinity between later Kondopoga individuals and modern Finnic groups (Figure 6D) is consistent with this interpretation and suggests that interactions with neighbouring Finnic populations may have contributed to the introduction of Eastern Eurasian-related ancestry into the Kondopoga gene pool. The absence of a significant signal for modern Finnish, Karelian, and Veps populations in this particular f4 test does not necessarily contradict this interpretation, as the Eastern Eurasian-related component in these populations is relatively minor and may be more effectively detected using other statistical approaches.

We next asked how much of the ancestry of each individual could be attributed to a source carrying Siberian-related ancestry, and whether that proportion changed along the pedigree. Rather than modelling the temporal groups, we fitted two-source qpAdm models to each of the 22 individuals separately, on 251,409 SNPs and against ten right populations (Materials and Methods). Four source pairs were tested, combining Estonia_IA.SG or Estonia_EarlyViking.SG as the first source with Finland_Levanluhta.AG or Russia_BolshoyOleniyOstrov_MBA as the second (Supplementary Materials, Table S14). The pair Estonia_EarlyViking.SG and Russia_BolshoyOleniyOstrov_MBA performed best: 19 of the 22 individuals satisfied all three criteria - a model not rejected at p > 0.05, both coefficients between zero and one, and each coefficient exceeding twice its standard error - with standard errors between 0.025 and 0.037, and only P290♀, P297♂ and P303♂ rejected (Figure 6B).

The estimated Siberian-related proportion separates the founding generations from everyone else buried at Kondopoga. It is 0.054 in P273♀, 0.076 in P274♂, 0.112 in P288♂ and 0.137 in P276♂, a group mean of 0.093 ± 0.014, against 0.191 ± 0.017 in their descendants P275♂, P282♂ and P285♀ and 0.214 ± 0.008 in the fifteen remaining individuals. The separation is complete: every member of the founding generations has a lower estimate than every other individual in the cemetery (Mann–Whitney p = 0.0005 for the founding generations against the late group). Replacing the second source with Finland_Levanluhta.AG raises the absolute values to 0.162, 0.352 and 0.393 for the three groups, because Levänluhta is itself admixed and a larger fraction of it is required to carry the same Siberian-related ancestry, but leaves the ordering and its significance unchanged. Using Estonia_IA.SG as the first source fails: the founding generations then receive a negative second-source coefficient (−0.022 ± 0.018), as expected if Iron Age Estonians already carry part of the ancestry being modelled.

Identity-by-descent also links the Kondopoga individuals to people buried at other sites. For the autosomal IBD analysis, we included selected samples that were closest to the Kondopoga individuals in time and geographic location. Raw sequencing data for samples from Andreeva et al. (2025) were kindly provided by the E. Rogaev group prior to their public release. Although we did not identify close relatives of the Kondopoga individuals among samples from other locations, IBD analysis revealed more distant relationships. P279, P294, and P295 shared IBD segments longer than 20 cM with individuals from medieval North Rus (Nefedyevo and Minino, present-day Vologda Oblast, Russia). All three successive male generations from 12th–13th century Kondopoga (P276 → P274 → P275) shared IBD segments >20 cM with an 11th-century male individual from Minino (present-day Vologda Oblast). All IBD connections >20 cM, including those with the northern Rus’ individuals, are shown in Supplementary Figures, Figure S2.

When IBD segments longer than 12 cM were considered, we identified additional connections between Kondopoga individuals and individuals from medieval South Finland, Viking-period individuals from the territory of present-day Russia and the Baltic region, including VK19 from Staraya Ladoga (Leningrad Oblast) and VK56 from Gotland, as well as medieval North Rus samples. Notably, P273, the female individual from the first Kondopoga couple, had no IBD segments with Finnish samples but shared IBD segments with Ya12 from Yaroslavl (North-Eastern Rus) (Supplementary Figures, Figure S10).

### Phenotypes

We determined eye, hair, and skin color for all 22 individuals (Supplementary Materials, Table S17). The majority showed the highest probabilities for blue eyes, pale to intermediate skin pigmentation, and blond to brown hair. Only three samples (P283, P297, and P298), which had fewer than 400,000 SNPs in the original data before imputation, showed higher probabilities for brown eyes. These results are broadly consistent with previous phenotypic reconstructions of medieval Rus populations, particularly those from Northern Rus, which indicate predominantly blue eyes, light hair, and pale to intermediate skin pigmentation (Andreeva et al., 2025). The phenotypes of the Kondopoga individuals appear largely homogeneous, with the exception of the first woman, P273, and her sons, P274 and P288, who had a 16–19% probability of red hair. The archaeological description confirms that P274 had light reddish hair and beard, while his son P275 had light-colored hair and beard (Archaeological Supplementary, Burial 356 and 358).

## DISCUSSION

Kondopoga is the first medieval cemetery in Karelia to be investigated using palaeogenetic methods and the first in the region for which a multigenerational pedigree has been reconstructed. Combining genetic relatedness, autosomal and X-chromosomal identity-by-descent (IBD), runs of homozygosity (ROH), uniparental markers and burial context allowed us to reconstruct the relationships between individuals buried at the site. Eight individuals form a pedigree spanning five generations, anchored by a couple buried among the earliest graves. IBD analysis extends the reconstructed relationships beyond the third degree: both founders share segments longer than 20 cM with 15 of the 20 other individuals, and 149 of the 231 pairs share IBD segments. The sampled burial community was therefore dominated by one extended family, although a second, shorter paternal line is represented by P298 and his son P300.

The genetic evidence points to a patrilineal family structure. The paternal lineage of the founding male recurs throughout the reconstructed pedigree, whereas mitochondrial lineages vary between individuals within the same generation. In one case, the mitochondrial lineage of P275 can be attributed to an unsampled mother. The pattern is consistent with patrilocal residence, in which women joined the community through marriage while men remained connected to the local kin group. X-chromosomal sharing provides further evidence for this interpretation. Male–male pairs do not share an X chromosome, whereas mother–son pairs share the expected X-chromosomal segments. These patterns also help distinguish lines of descent through sons from those through daughters. No direct female descendants of the founding couple were identified among the sampled individuals from the earliest generations, consistent with the possibility that daughters married into other communities and were buried elsewhere. This interpretation remains tentative because the cemetery provides only a partial record of the family.

ROH results suggest that this patrilineal system did not depend on close-relative unions. None of the analysed individuals carries a ROH longer than 20 cM, indicating an absence of evidence for recent close inbreeding. Partners could therefore have come from outside the immediate kin group even as descendants of the founding couple continued to use the same burial ground. Comparable combinations of patrilineal descent, patrilocal residence and female exogamy have been reconstructed at Bronze Age and early medieval sites in central Europe (Mittnik et al., 2019; Gnecchi-Ruscone et al., 2024). Kondopoga provides a northern example of a similar pattern, documented across several centuries in a medieval cemetery in the Novgorod-connected lands.

The earliest burials suggest that the founding couple held a distinctive position within the community. Their graves occupy the highest, central part of the hill, and P273 is buried in the site’s only stone cist, accompanied by gold and silver ornaments.Their genetic profiles also differ from those of later individuals. P273 and her sons occupy a distinct position in both the Eurasia-wide and regional principal component analyses and show the lowest proportions of Uralic-related ancestry in the sampled group. Individual-level f3 analyses place their strongest affinities among Baltic- and Slavic-associated groups rather than Finnic populations or individuals from the northern medieval Rus’ cemeteries included in the comparison. P273 also has the lowest background relatedness in the dataset, with no ROH longer than 2.7 cM. Together, the burial context and genetic results suggest that the founders came from a population with a different ancestry profile and mating history from those of later generations. Their graves mark the beginning of the cemetery, which their descendants continued to use.

The persistence of the paternal lineage is especially clear when Y-chromosomal and autosomal evidence are considered together. The founding male’s Y-chromosomal lineage, R-YP418*, is present in P276 in the twelfth–thirteenth centuries, his son P288, his fourth-generation descendant P282, and P303, whose burial dates to no later than the mid-sixteenth century. P294, dated to the fifteenth–sixteenth centuries, carries the ancestral branch R-YP417*. P297, dated to the sixteenth–seventeenth centuries, carries R-PF6155*, an upstream position within the same broader R1a lineage. This is compatible with the same paternal origin but does not establish it.

Autosomal relatedness, by contrast, decreases with each generation as DNA is recombined and passed down from multiple ancestors. P303 shares 227 cM with P275, 155 cM with P288 and 141 cM with each of P274 and P276, consistent with more distant relationships than those seen among the earlier generations. His closest relative in the dataset is P278, at the third degree. The contrast between the persistence of a Y-chromosomal lineage and declining autosomal sharing illustrates why the two types of evidence are complementary: together, they allow paternal continuity to be traced even when autosomal relatedness to the founders has become more distant.

One individual remains difficult to place within this chronology. P297 is assigned third-degree relationships to P274 and P275 based on 813 cM and 534 cM of sharing, respectively, including a segment of 99 cM. These estimates are difficult to reconcile with the sixteenth–seventeenth-century date of his burial. Neither background relatedness, which would be expected to produce shorter segments, nor the available contamination estimates provides a clear explanation. We therefore retain the discrepancy rather than propose a genealogical arrangement unsupported by the current evidence.

The ancestry of the family also changed across generations. The founding couple and their sons have the lowest proportions of Uralic-related ancestry in the cemetery. This proportion rises in their grandson P275 to levels observed among other members of the burial community and is higher still in P282, a fourth-generation descendant. It remains at a similar level among the latest individuals. The father–son pair P274 and P275 is particularly informative because their positions in the pedigree are established. The increase in Uralic-related ancestry between them must reflect ancestry contributed by P275’s mother, who is not sampled directly. His mitochondrial haplogroup, H1n4, also differs from that of the founding female.

These findings suggest that women marrying into the family contributed ancestry that differed from that of the founders. The f4 statistics support this interpretation, showing increased relative affinity to Eastern Eurasian- and Siberian-related populations among later individuals. Individual-level admixture models estimate a rise in Siberian-related ancestry from 0.093 ± 0.014 in the founding generations to 0.191 ± 0.017 among their descendants, with no overlap between the two sets of individuals. The combination of paternal continuity, changing mitochondrial lineages and increasing Siberian-related ancestry is consistent with repeated incorporation of women from populations with different ancestry profiles, rather than replacement of the family by a new group.

The regional context helps narrow down, but not identify, the source of this ancestry. Modern Ludic Karelians inhabit parts of southeastern Karelia, including the Kondopoga area, while Veps communities are found across Karelia and neighbouring regions around Lake Onega. Both are geographically plausible sources of partners for the family. Outgroup-f3 results show the strongest affinities among Veps, Ingrian Finns, and Livvi or Ludic Karelians in the later groups. However, these populations are genetically similar enough that the two-source models cannot reliably distinguish between them. Moreover, medieval communities may have differed from their present-day descendants.

The admixture models should therefore be interpreted as estimates of ancestry components, not as direct identifications of the people who contributed them. Russia_BolshoyOleniyOstrov_MBA, a Bronze Age population from the Kola Peninsula, serves as a proxy for Siberian-related ancestry because it carries this component in a comparatively unadmixed form. It does not represent a demonstrated source population for the Kondopoga family. Similarly, the ancestry proportions do not establish the language, cultural identity or ethnicity of individual women who married into the family.

The cemetery’s long period of use also makes it possible to compare the genetic pedigree with the spatial arrangement of the graves. The founding couple, their sons and their grandson occupy a compact area in the highest central part of the hill, while later descendants were buried around them. This correspondence suggests that biological relationships were reflected, at least in part, in the organisation of the burial ground. The genetic continuity of the family was thus accompanied by continuity in its use of a particular place within the cemetery.

The ancestry of the founders raises a broader question about population movements in the lands connected to Novgorod. Craniological studies have reported that eleventh–thirteenth-century series from the Novgorod lands resemble medieval and modern Baltic populations more closely than later series, which show greater similarity to Finnish populations. These changes have been attributed to admixture during the medieval period (Alekseeva and Buzhilova, 1997). Baltic morphological features have also been described in medieval Karelian series from the northern Ladoga region and in the stone graves at Raglitsy and Kylalahti Kalmistomäki (Khartanovich and Shirobokov, 2010; Sankina, 2000, 2008). The genetic affinities of the earliest Kondopoga individuals are consistent with this broader pattern. IBD segments also connect members of the family to medieval individuals from northwestern Rus’, southern Finland and the Baltic region, including VK19 from Staraya Ladoga and VK56 from Gotland. These segments do not indicate close genealogical relationships, however, and cannot establish a direct connection between the founders and any particular site. Whether the family’s arrival at Kondopoga formed part of a wider movement of people associated with the Novgorod lands remains an open question. Testing this hypothesis, which fits general historical data, will require comparable pedigrees from additional medieval cemeteries in Karelia and the Novgorod lands.

Kondopoga provides the first multigenerational pedigree reconstructed from a medieval cemetery in Karelia. The results show that one family used the burial ground over several centuries, maintaining a paternal lineage while its autosomal ancestry changed across generations. The contrast between stable Y-chromosomal lineages and changing mitochondrial and autosomal profiles points to a sustained contribution from women who married into the family. By combining genetic relatedness with burial context, this study reconstructs aspects of a medieval family’s history that would be difficult to recover from either genetic or archaeological evidence alone. Comparable studies across the region will help establish how often this pattern occurred and how it related to wider population changes in the lands connected to Novgorod.

## MATERIALS AND METHODS

All procedures involving ancient DNA (aDNA) were carried out in a dedicated clean-room environment at the Federal Research Center of Biotechnology of the Russian Academy of Sciences (K. G. Skryabin Institute of Bioengineering). DNA extraction from teeth, bones, auditory ossicles, and petrous bones was performed using a magnetic separation approach with buffer D following the published protocol of Rohland et al. (2018; doi:10.1038/s41596-018-0050-5). Extracted DNA was subsequently converted into single-stranded libraries using the xGen™ ssDNA & Low-Input DNA Library Preparation Kit (IDT, USA), following the published protocol of Gansauge and Meyer (2013; doi:10.1038/nprot.2013.038). In-solution enrichment for target genomic regions was carried out with the MyBaits Expert Human Affinities Prime Plus Kit (Daicel Arbor Biosciences). Sequencing was performed on an Illumina HiSeq 1500 platform (Illumina, USA) in paired-end format (2 × 150 bp). An initial test sequencing run was conducted on the constructed libraries to assess low-coverage DNA fragments, generating approximately 3–4 million single-end reads (50 bp in length) per sample. Libraries with less than 1% endogenous DNA at this test sequencing stage were not submitted to capture enrichment and are not considered further in this study. For samples showing the highest levels of endogeneity and characteristic C > T substitutions at the 5′ ends of DNA fragments, an additional library was prepared from the same DNA extract following pretreatment with a mixture of uracil-DNA glycosylase (UDG) and endonuclease VIII (Gansauge and Meyer, 2013).

The study included 22 individuals (10 female and 12 male, as determined from the genetic data) from the Kondopoga necropolis, selected based on archaeological context and the availability and preservation of skeletal material for genetic analysis. The estimated age at death, determined using anthropological methods, ranged from under 1 year old to over 50 years old and is provided in Supplementary Table S1 and the Archaeological Supplementary Material.

Archaeological fieldwork at the Kondopoga necropolis was conducted under permission (Open Sheet) No. R-18-00103-00/01106204, issued on 28 March 2024. Samples were included in downstream analyses according to the coverage and data-quality requirements specified for each method. Consequently, the number of individuals varied among analyses. The sample size was determined by the number of suitable archaeological specimens available and the amount and preservation of endogenous DNA recovered from the samples. No formal a priori power analysis was performed because the study was based on a finite archaeological sample that could not be increased experimentally. Randomization was not applicable because the study did not involve experimental treatment groups or randomized allocation of samples. To reduce potential analytical bias, genetic and computational analyses were performed using predefined DNA library identifiers, without access to the corresponding archaeological identities of the samples during the initial analysis. The analyst was therefore unaware of the individual archaeological context associated with each library identifier until the primary analytical results had been obtained.

Raw sequencing data were processed using the nf-core/eager pipeline v.2.5.3 (https://nf-co.re/eager) (Yates et al., 2021; Peltzer et al., 2016) including the tools described below. Adapter trimming and merging of paired-end reads were performed with AdapterRemoval (RRID:SCR_011834) v.2.3.2 (Schubert et al., 2016) using default parameters, and only merged reads were retained for downstream analysis. Reads were mapped to the human reference genome (GRCh37) with Bowtie 2 (RRID:SCR_016368) v.2.4.4 (Langmead and Salzberg, 2012) applying settings optimised for ancient DNA (--very-sensitive-local, -k 100). Duplicate reads were removed with DeDup v.0.12.8 (Peltzer et al., 2016). To eliminate residual cytosine deamination, the first and last two bases of each read were trimmed using the trimBam module of bamUtil v.1.0.15 (Jun et al., 2015). Postmortem damage patterns were estimated on a subset of 10,000 reads with mapDamage (RRID:SCR_001240) v.2.2.1 (Jónsson et al., 2013). Alignments were further filtered to retain only reads with a mapping quality ≥20 and a minimum read length of 30 bp using SAMTOOLS (RRID:SCR_002105) v.1.12 (Li et al., 2009).

Genotyping was performed on trimmed BAM files using pileupCaller from sequenceTools v.1.5.2 (https://github.com/stschiff/sequenceTools) with the 1240K SNP panel. Random pseudo-haploid calls were generated at each target position using the --randomHaploid option. Genetic sex was determined for all individuals based on the ratio of reads mapped to the X and Y chromosomes (Skoglund et al., 2013). Nuclear contamination was assessed on the X chromosome using ANGSD (RRID:SCR_021865) v.0.935 (Korneliussen et al., 2014) with parameters -doCounts 1 -doMaf 1 -doGlf 2 -doMajorMinor 1 and restricting to males. The mitochondrial-to-nuclear DNA ratio was estimated with MTNucRatioCalculator v.0.7.1 (https://github.com/apeltzer/MTNucRatioCalculator).

Mitochondrial contamination levels were estimated using Schmutzi v.1.5.7 (Renaud et al., 2015). Mitochondrial haplogroups were determined with Haplogrep3 v.3.2.1 (Schönherr et al., 2023) based on the PhyloTree.org (RRID:SCR_012948) build 17. Y-chromosome haplogroups were determined from BAM files using Yleaf v3.2.1 (Ralf et al., 2018). Biological relatedness up to the third degree was investigated using READv2 (RRID:SCR_026959) v.2.01 (Alaçamlı et al., 2024). READv2 was run on pseudo-haploid genotype data with default parameters, enabling the classification of first-, second-, and third-degree relatives, as well as the differentiation between parent–offspring and full-sibling relationships. To assess possible parental relatedness, we calculated runs of homozygosity (ROHs) using hapROH (Ringbauer et al., 2021) on the pseudo-haploid genotype data. For the ROH analysis, three of the 22 Kondopoga individuals were excluded because they had fewer than 400,000 SNPs, leaving 19 individuals for this analysis. The inferred ROHs were further used to estimate the effective population size.

To explore the genetic affinities of the ancient individuals, we performed principal component analysis (PCA) using the smartpca tool from the Eigensoft (RRID:SCR_004965) v.8.0.0 package (Price et al., 2006; Patterson et al., 2006). We computed principal components using a reference panel of 79 modern populations from the Human Origins (HO) dataset. Principal components were then used to project the ancient individuals onto the same genetic space. Published comparative datasets used in this study were obtained from the Allen Ancient DNA Resource (AADR) v.62 (Mallick et al., 2024), and from the datasets reported by Gretzinger et al. (2025) (ENA accession PRJEB81250), Nordfors et al. (2025) (ENA accession PRJEB88970), and Agdzhoyan et al. (2024), available through the Russian Gene Pool project database. Projection was performed using the lsqproject: YES option in smartpca.

To investigate the ancestry composition of the Kondopoga samples in the context of modern populations and other ancient individuals from relevant temporal and geographic backgrounds, we performed ADMIXTURE (RRID:SCR_001263) v.1.3.0 analysis (Alexander et al., 2009). The analysis was run for a range of ancestral components (K) from 6 to 12, and the cross-validation (CV) error was used to select the most plausible number of components. The lowest CV error was observed at K = 12 (Supplementary Materials, Table S8). For visualization, we focused on K=6, which provided a relatively simple and interpretable representation of the inferred ancestry components. The input dataset consisted of 13,931 samples (including both ancient and modern individuals, Supplementary Materials, Table S7) with more than 30,000 SNPs overlapping the 1240K panel of the Allen Ancient DNA Resource (AADR) (Mallick et al., 2024). Prior to ADMIXTURE, we applied quality control and linkage disequilibrium (LD) pruning using PLINK (RRID:SCR_001757) v.1.9.0-b.8 (Purcell et al., 2007). First, markers with minor allele frequency (MAF) below 0.01 were removed. LD pruning was performed using the --indep-pairwise option with a window size of 50 SNPs, a step size of 2 SNPs, and an r² threshold of 0.5. To test hypotheses of population admixture, we calculated f3-outgroup and f4 statistics using the qp3Pop and qpDstat programs from the ADMIXTOOLS (RRID:SCR_018495) v.8.0.2 package (Patterson et al., 2012). For outgroup f3 statistics, we used the inbreed: YES option. Standard errors for f3 and f4 statistics were estimated using ADMIXTOOLS. Two-source admixture models were fitted with qpAdm as implemented in ADMIXTOOLS 2 v.2.0.10 (Maier et al., 2023), using 251,409 autosomal SNPs. Each of the 22 Kondopoga individuals was modelled separately. The right (reference) set comprised Mbuti.HO, Yoruba.HO, Han.HO, Papuan.HO, Nganasan.HO, Russia_YuzhniyOleniyOstrov_Mesolithic.AG, Luxembourg_Mesolithic.DG, Austria_N_LBK.AG, Georgia_Kotias_Mesolithic.SG and Russia_MA1_UP.SG, chosen to span the African, East Asian, Oceanian, Siberian, eastern and western hunter-gatherer, early farmer, Caucasus hunter-gatherer and Ancient North Eurasian axes. Four pairs of left-hand sources were tested, combining the Iron Age Estonian group (Estonia_IA.SG) or the Early Viking Age Estonian group (Estonia_EarlyViking.SG) with the Levänluhta group (Finland_Levanluhta.AG) or the Bolshoy Oleniy Ostrov group (Russia_BolshoyOleniyOstrov_MBA); for the last of these, the .AG and .SG samples were pooled. A model was considered admissible when the rank-1 p-value exceeded 0.05, both coefficients lay between zero and one, and each coefficient exceeded twice its standard error, the last criterion testing whether the two-source model is supported over a single-source model. Standard errors were obtained by block jackknife over 5-cM blocks. Full results for all 88 models are given in Supplementary Materials, Table S14.

To investigate distant kinship relationships beyond the third degree between the Kondopoga samples and other selected ancient individuals, we performed IBD analysis using ancIBD v.0.5 (Ringbauer et al., 2024), which requires phased and imputed genotype data as input.

Genotype imputation was performed with GLIMPSE2 v.2.0.0 (Rubinacci et al., 2023; Rubinacci et al., 2021) using the 1000 Genomes Project reference panel (RRID:SCR_008801). All 22 Kondopoga individuals were retained for the IBD analysis, as all met the minimum coverage threshold of 298,000 SNPs on the 1240K panel. Other ancient samples were included only if they had more than 600,000 SNPs covered on the same panel. IBD segments were called using minimum length thresholds of 8 cM, 12 cM, 16 cM, and 20 cM.We constructed an IBD network and visualized it using the MultiGravity ForceAtlas2 (Jacomy et al., 2014) force-directed layout algorithm, with clusters identified using the Leiden algorithm (Traag et al., 2019).

Phenotypic traits (eye, hair, and skin colour) were predicted using the HIrisPlex system (RRID:SCR_014058) (Walsh et al., 2014), which relies on a panel of 41 genetic markers. Because the original genotyping data did not cover all required single-nucleotide polymorphisms (SNPs), we used the imputed VCF files generated during the imputation step described above. Recent work has shown that for ancient genomes with coverage in the range of 0.1–0.5×, imputation yields phenotype predictions that are comparable to those obtained directly from genotype data (Maróti et al., 2026).

The chronologies used in this study were established using archaeological methods, including stratigraphy, typology, planigraphy, and analysis of associated grave goods. No radiocarbon dating was performed. Descriptions of the burials and associated grave goods are provided in the Supplementary Material.

### Pedigree reconstruction

The pedigree was assembled from five kinds of evidence. Pairwise coefficients from READv2 defined the first-, second- and third-degree links. The direction of descent was fixed independently of the genetic data, from the archaeological dating of the graves and the age at death recorded for each individual, and the reconstruction was anchored on the earliest burials. Mitochondrial and Y-chromosomal haplogroups were used as consistency checks, so that a mother–child or father–son link was accepted only where the corresponding uniparental lineage was shared. Where two genealogical configurations of the same autosomal degree were possible, X-chromosomal identity-by-descent was used to distinguish between them and to establish the sex of the unsampled individual through whom a connection passes, on the expectation that a man transmits his single X chromosome to each of his daughters and none to his sons. Identity-by-descent segments called with ancIBD then extended the reconstruction to fourth- to sixth-degree pairs; those whose position in the pedigree could not be fixed are reported as relatives of the kindred without a placement, and the one case in which the genetic and the archaeological evidence conflict is reported as unresolved rather than resolved by assumption.

## DATA AVAILABILITY

The data for this study have been deposited in the European Nucleotide Archive (ENA) at EMBL-EBI under accession number PRJEB128067 (https://www.ebi.ac.uk/ena/browser/view/PRJEB128067). The data will be made publicly available upon publication of the article as a reviewed preprint, and are available to editors and reviewers on request during assessment.

## ACKNOWLEDGEMENTS

We thank E. I. Rogaev and his group for providing raw sequencing data for the individuals reported in Andreeva et al. (2025) prior to their public release, which made the comparison of the Kondopoga individuals with medieval populations of northern Rus’ possible. We thank the Institute of Archaeology of the Russian Academy of Sciences for access to the archaeological material from the Kondopoga necropolis and for the photographs reproduced in Figure 1 and Archaeological Supplementary.

## AUTHOR CONTRIBUTIONS

D.A.V. — conceptualization, methodology, data curation, formal analysis, visualization, writing – original draft. M.V.L. — investigation, writing – review and editing. K.V.Z. — investigation, writing – review and editing. I.K.R. — investigation, resources, writing – original draft (archaeological supplementary). A.V.N. — investigation, resources, writing – original draft (archaeological supplementary). M.V.D. — investigation, resources, writing – review and editing. N.A.M. — funding acquisition, investigation, project administration, resources, writing – review and editing. E.B.P. — conceptualization, resources, funding acquisition, project administration, supervision, writing – review and editing.

## ETHICS AND PERMITS

Archaeological fieldwork at the Kondopoga necropolis was conducted under permission (Open Sheet) No. R-18-00103-00/01106204, issued on 28 March 2024 by the Ministry of Culture of the Russian Federation, and the skeletal material is curated by the Institute of Archaeology of the Russian Academy of Sciences.

